# Conditional Analysis Cakewalk

**DOI:** 10.1101/2020.02.14.947978

**Authors:** Jennifer N. French, Brady J. Gaynor, James A. Perry

## Abstract

**Summary:** Sequential conditional analysis is an important tool for identifying the unique signals in single-variant association analysis between genomic variants and various phenotypic outcomes. With large datasets, it is tedious and time-consuming to manually run the sequential steps which require the association analysis, computation of linkage disequilibrium (LD) between the variants, generation of a visualization graphic, and identification of the “top hit” to be inserted as an additional covariate for the next iteration. The Conditional Analysis Cakewalk automates this process, making it easy and fast, while only requiring user input in the initial configuration steps.

**Availability and Implementation:** The Conditional Analysis Cakewalk is implemented as a Linux shell script utilizing MMAP or PLINK and LocusZoom. Source code and documentation are available at https://github.com/jennetics/Cakewalk.

## 1. Introduction

Genetic association studies have been enormously helpful in identifying loci that map to numerous traits and diseases. Frequently, the associated loci include multiple associated variants that are highly correlated (i.e., in high linkage disequilibrium (LD)). Conditional association analysis is often used as a next step to identify the variant or variants at the associated loci that are independently associated with the trait or disease. However, sequential conditional analysis can be tedious and time-consuming, especially when there are numerous variants that are highly correlated. To address this problem, we have developed the Conditional Analysis Cakewalk tool for automating this important process.

## 2. Implementation

### 2.1 Inputting variables

The first step in the Conditional Analysis Cakewalk is to provide user-specified inputs. These include the input data file, trait of interest, chromosome number, start/end positions for the region to be scanned, as well as covariates and the desired suffix for output files. The user has the option to have the “top hit” automatically selected for them. If this option is not selected, a list will print for manual selection of the reference variant (Supplementary Figure 1). Additionally, there are four thresholds that can be adjusted: maximum p-values for including variants in both the initial and subsequent LD calculations; the p-value for establishing significance of variants as the termination condition, and a p-value for limiting which variants are included in each association analysis. The final p-value was included to improve processing times. The y-axis range and significance lines within the LocusZoom plot can also be specified. Finally, the results will be routed to a user-specified directory with an option for removing all other files to conserve storage space.

### 2.2 Association analysis, LD calculation and visualization

The Conditional Analysis Cakewalk tool is available in two scripts: one utilizing MMAP and the other utilizing PLINK. For association analysis on related subjects, MMAP provides a mixed model approach (O’Connell). PLINK can be used when relatedness does not need to be included in the analysis (Purcell et al., 1007). In both scripts, the association output file is sorted and processed into a marker list for LD calculations, which are performed with either the MMAP or PLINK program. The LD results are again sorted and processed into files formatted specifically for LocusZoom (Welch & Pruim, 2010). The LocusZoom plot includes all variants in the specified region. There will be a green vertical line on the plot in the location of the top variant. A boxplot will be generated for each top variant, as well as a final plot with allele counts for all top variants combined, using R (R Core Team, 2013). The Conditional Analysis Cakewalk will continue to iterate through the association analyses, LD calculations, boxplot and LocusZoom plotting until the p-value of the top variant in an iteration is greater than the specified cutoff p-value.

### 2.3 The results directory

All results will be placed in a user-specified results directory. This directory will contain the LocusZoom plot and boxplot for each step of the conditional analysis, as well as a file entitled “TOP_SNPS.csv”. This file will contain the variant name, p-value, β, and file name for the LocusZoom and boxplot for every step of the analysis.

## 3. Example on simulated data

As an example, we performed a conditional sequential association analysis of low-density cholesterol (LDL-C) levels within the LDLR gene using the Conditional Analysis Cakewalk with simulated data. None of the variants identified in this simulated example should be assumed to have any clinical or other significance. The analysis included 404 subjects and 117,982 variants. The association analysis was conducted with MMAP using chromosome 19, region 8,000,000-18,000,000, which contains the LDLR gene, a gene shown to have a connection with LDL levels (Bjune, Sundvold, Leren, & Naderi, 2018; Tang et al., 2018; Yang et al., 2015). The initial association analysis identified 212 variants within this region as significantly associated with LDL (p ≤ 10^−8^). 12 of these variants were found to be in LD with the top variant, chr19:11209757 (p = 1.50 x 10^−198^; r^2^ ≥ 0.80; Figure 1). After adjusting for chr19:11209757 by adding it as a covariant to the analysis model, variant chr19:11242044 was found to be significantly associated with LDL (p = 1.90 x 10^−67^). The second association analysis found 123 variants significantly associated with LDL (p ≤ 10^−8^), 45 of which were in LD with chr19:11242044 (r^2^ ≥ 0.80; Figure 2). Continuing the conditional analysis, chr19:11242044 was added to the association analysis as a covariate, and no additional variants were found to be independently associated with LDL in the specified region. Boxplots for each iteration individually and combined can be found in the supplementary data.

**Figure 1.**
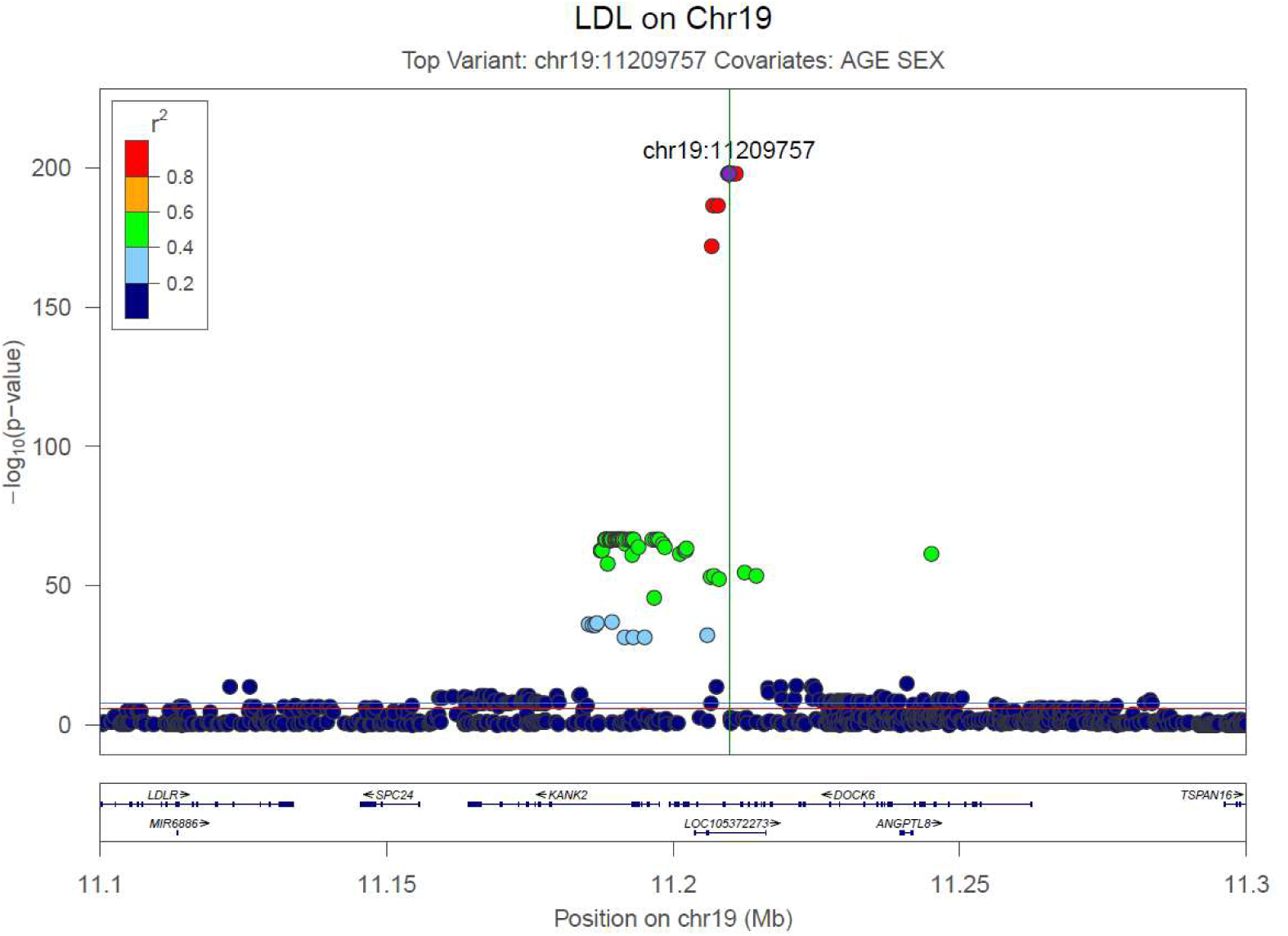
LocusZoom plot for 19:11086210:T:C. This LocusZoom plot is from the first iteration of the Conditional Analysis Cakewalk using simulated data. It displays SNPs in LD with the top SNP.

**Figure 2.**
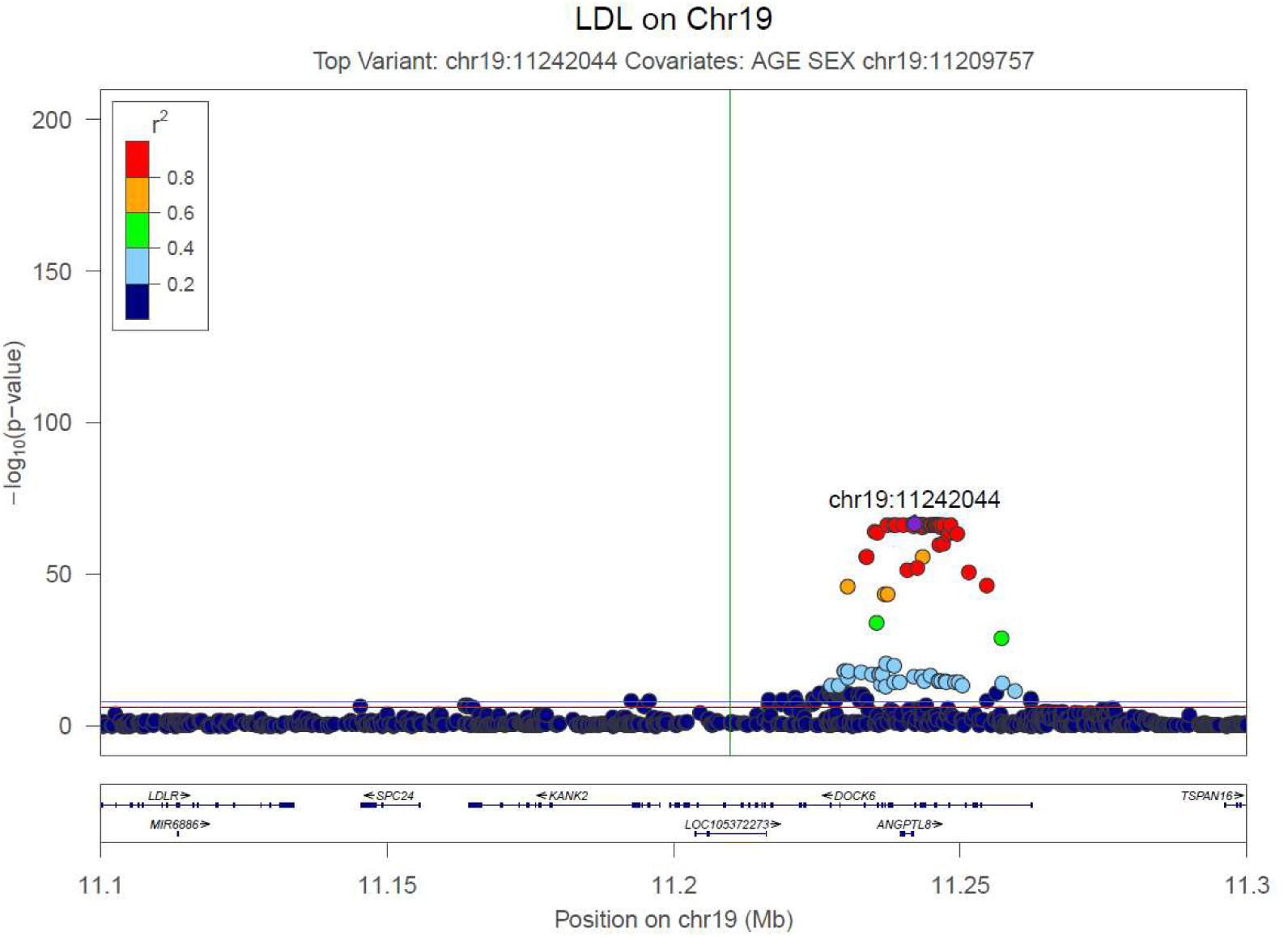
LocusZoom plot for 19:11156017:T:G. This LocusZoom plot is from the second iteration of the Conditional Analysis Cakewalk using simulated data. It displays SNPs in LD with the top SNP after adjusting for 19:11086210:T:C.

## 4. Limitations

A limitation of the Conditional Analysis Cakewalk is that it may select a single variant arbitrarily from within a group of near perfectly correlated variants, and not necessarily the variant that is most likely to be “causal,” unlike Bayesian methods which allow for adding weights to variants, such as CAVIARBG, BIMBAM, and PAINTOR (Chen, et al., 2015). To address this limitation, the user has the option to manually select the top variant for analysis.

## Acknowledgements

The authors acknowledge the developers of MMAP, Plink and LocusZoom.

## References

Bjune, K., Sundvold, H., Leren, T.P., & Naderi, S. (2018). MK-2206, an allosteric inhibitor of AKT, stimulates LDLR expression and LDL uptake: A potential hypocholesterolemic agent. Atherosclerosis, 276, 28–38.

Chen, W., Larabee, B.R., Ovsyannikova, I.G., Kennedy, R.B., Haralambieva, I.H., Poland, G.A., & Schaid, D.J. (2015). Fine mapping causal variants with an approximate Bayesian method using marginal test statistics. Genetics, 200 (3), 719–736. Doi: 10.1534/genetics.115.176107.

O’Connell, J.R. MMAP: Mixed model analysis for pedigrees and populations. Available from: https://mmap.github.io/.

Purcell, S, Neale, B, Todd-Brown, K, Thomas, L, Ferreira, MAR, Bender, D, Maller, J, Sklar, P, de Bakker, PIW, Daly, MJ, Sham PC. (2007). PLINK: a toolset for whole-genome association and population-based linkage analysis. American Journal of Human Genetics, 81. Available from https://www.cog-genomics.org/plink/1.9/#test_note.

R Core Team. (2013). R: a language and environment for statistical computing. R Foundation for Statistical Computing, Vienna, Austria. http;//www.R-project.org/.

Tang, L., Wang, G., Jiang, L., Chen, P., Wang, W., Chen, J., & Wang, L. (2018). Rose of sEH R287Q in LDLR expression, LDL binding to LDLR and LDL internalization in BEL-7402 cells. Gene, 667, 95–100.

Welch, R., Pruim, R. (2010). LocusZoom standalone. Available from: https://github.com/statgen/locuszoom-standalone.

Yang, J.H., Bang, M.A., Jang, C.H., Jo, G.H., Jung, S.K., & Ki, S.H. (2015). Alginate oligosaccharide enhances LDL uptake via regulation of LDLR and PCSK9 expression. Journal of Nutritional Biochemistry, 26 (11), 1393–1400.

