## Supplementary Figures for "Conditional Analysis Cakewalk"

```
Enter the line number of the desired variant to select as the
reference variant (columns are SNP, p-value, beta):
     1  rs141820146  1.498e-198   175.2
     2  rs142444132  1.498e-198   175.2
     3   rs17248769  1.498e-198   175.2
     4   rs17248776  1.498e-198   175.2
     5   rs17248783  1.498e-198   175.2
     6    rs2228671  1.498e-198   175.2
     7   rs73015033  1.498e-198   175.2
     8   rs73015034  1.498e-198   175.2
     9   rs74857287  1.498e-198   175.2
    10  rs117423069  2.333e-187   173.2
    11   rs17242395  2.333e-187   173.2
```

**Supplementary Figure 1.** Sample of output when choosing to manually select reference variant.

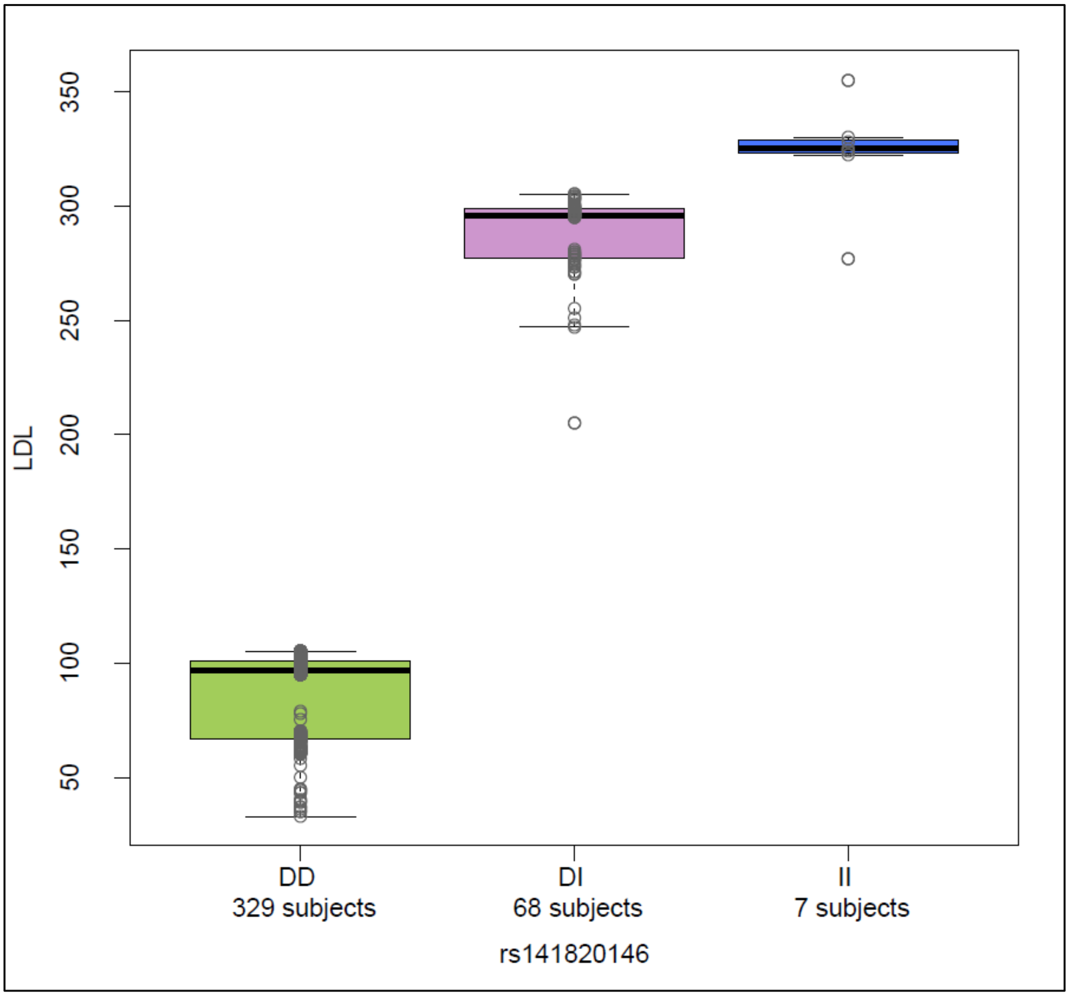


**Supplementary Figure 2.** Boxplot of LDL levels based on genotype of top variant from the first iteration of the Conditional Analysis Cakewalk on simulated data.

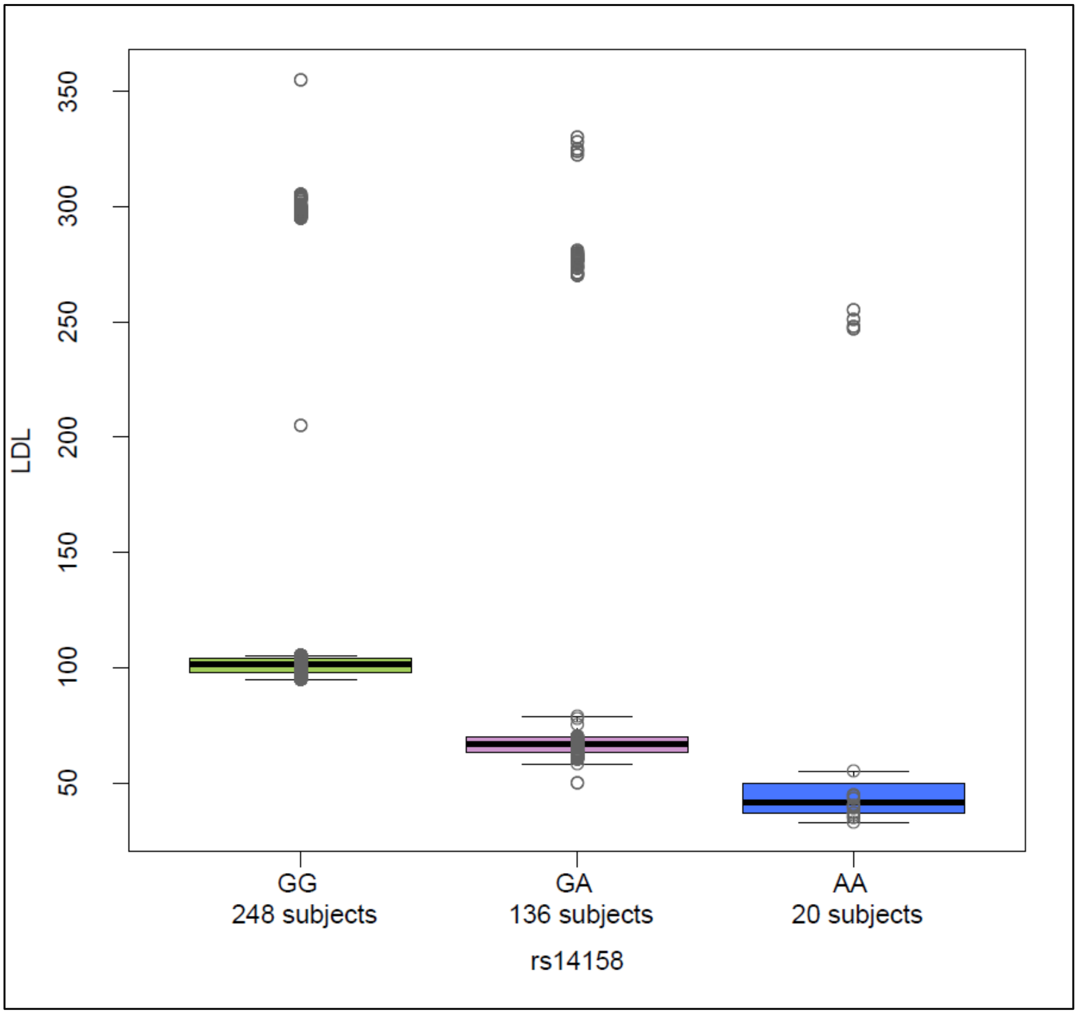


**Supplementary Figure 3.** Boxplot of LDL levels based on genotype of top variant from second iteration of Conditional Analysis Cakewalk on simulated data.

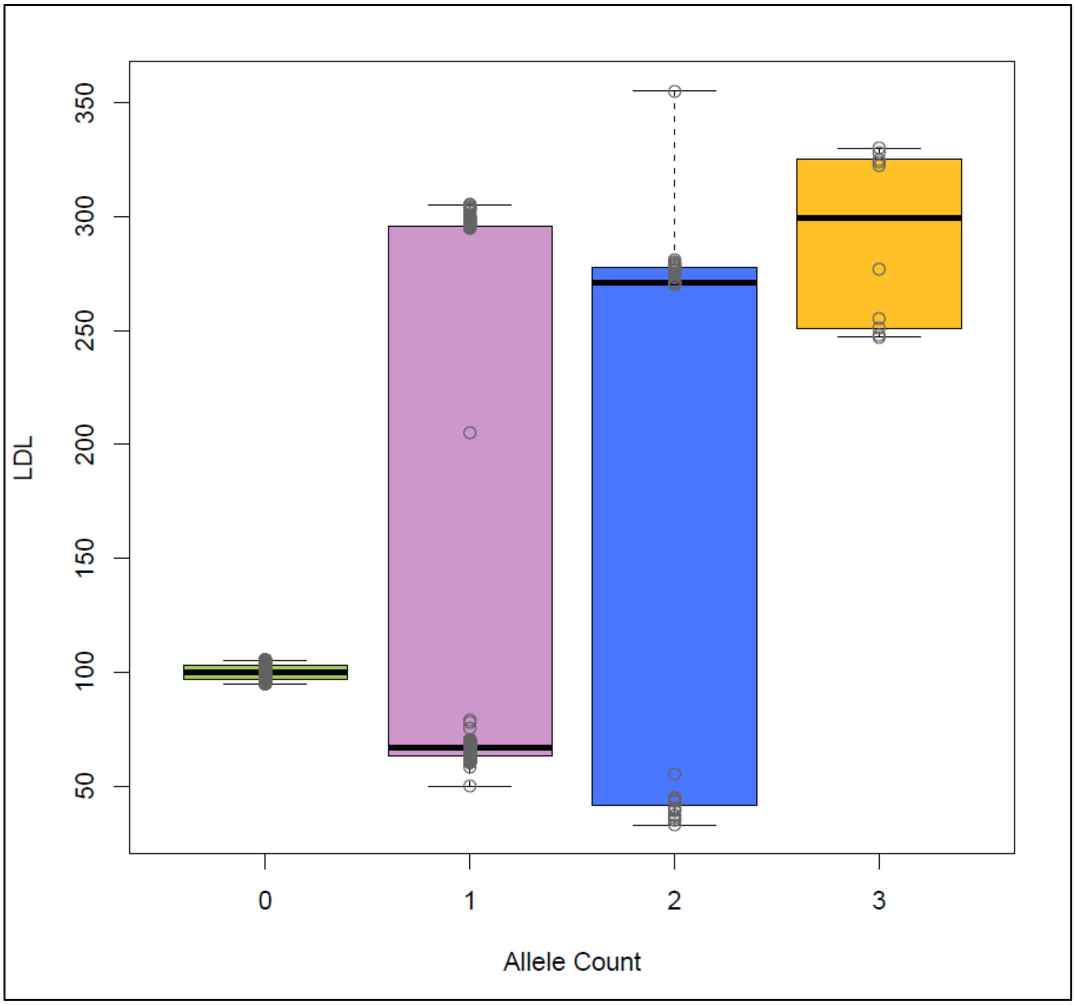


**Supplementary Figure 4.** Boxplot of LDL levels based on allele count of top variants from all iterations of the Conditional Analysis Cakewalk on simulated data.

| SNPNAME | CHR:POS | PVALUE | BETA | LOCUSZOOM | BOXPLOT |
|---|---|---|---|---|---|
| rs141820146 | chr19:11209757 | 1.50E-198 | 175.2 | LZ_LDL.chr19:11209757.pdf | LDL.test1.pdf |
| rs14158 | chr19:11242044 | 1.90E-67 | -32.9 | LZ_LDL.chr19:11242044.pdf | LDL.test2.pdf |

Statistics for final boxplot

| Allele_count | Subject_count |
|---|---|
| 0 | 205 |
| 1 | 150 |
| 2 | 39 |
| 3 | 10 |

**Supplementary Figure 5.** Contents of output file, TOP_SNPS.csv, containing data from each iteration of the Conditional Analysis Cakewalk on simulated data and statistics for the final boxplot.
